# Assessing Tumor Microenvironment Characteristics and Stratifying EPR with a Nanobubble Companion Nanoparticle via Contrast-Enhanced Ultrasound Imaging

**DOI:** 10.1101/2023.11.20.567934

**Authors:** Michaela B. Cooley, Dana Wegierak, Reshani Perera, Eric C. Abenojar, Pinunta A. Nittayacharn, Felipe M. Berg, Youjoung Kim, Michael C. Kolios, Agata A. Exner

**Affiliations:** Department of Biomedical Engineering, Case Western Reserve University, Cleveland, Ohio 44106, United States; Department of Radiology, Case Western Reserve University, Cleveland, Ohio 44106, United States; Department of Biomedical Engineering, Faculty of Engineering, Mahidol University, Puttamonthon, Nakorn Pathom, 73170, Thailand; Hospital Israelita Albert Einstein, São Paulo, São Paulo 05652-900, Brazil; Department of Physics, Toronto Metropolitan University, Toronto, Ontario, M5B 2K3, Canada

**Keywords:** nanobubbles, ultrasound contrast agents, companion nanoparticles, oncology, EPR

## Abstract

The tumor microenvironment is characterized by dysfunctional endothelial cells, resulting in heightened vascular permeability. Many nanoparticle-based drug delivery systems attempt to use this enhanced permeability combined with impaired lymphatic drainage (a concept known as the ‘enhanced permeability and retention effect’ or EPR effect) as the primary strategy for drug delivery, but this has not proven to be as clinically effective as anticipated. The specific mechanisms behind the inconsistent clinical outcomes of nanotherapeutics have not been clearly articulated, and the field has been hampered by a lack of accessible tools to study EPR-associated phenomena in clinically relevant scenarios. While medical imaging has tremendous potential to contribute to this area, it has not been broadly explored. This work examines, for the first time, the use of multiparametric dynamic contrast-enhanced ultrasound (CEUS) with a novel nanoscale contrast agent to examine tumor microenvironment characteristics noninvasively and in real-time. We demonstrate that CEUS imaging can: (1) evaluate tumor microenvironment features and (2) be used to help predict the distribution of doxorubicin-loaded liposomes in the tumor parenchyma. CEUS using nanobubbles (NBs) was carried out in two tumor types of high (LS174T) and low (U87) vascular permeability, and time-intensity curve (TIC) parameters were evaluated in both models prior to injection of doxorubicin liposomes. Consistently, LS174T tumors showed significantly different TIC parameters, including area under the rising curve (2.7x), time to peak intensity (1.9x) and decorrelation time (DT, 1.9x) compared to U87 tumors. Importantly, the DT parameter successfully predicted tumoral nanoparticle distribution (r = 0.86 ± 0.13). Ultimately, substantial differences in NB-CEUS generated parameters between LS174T and U87 tumors suggest that this method may be useful in determining tumor vascular permeability and could be used as a biomarker for identifying tumor characteristics and predicting sensitivity to nanoparticle-based therapies. These findings could ultimately be applied to predicting treatment efficacy and to evaluating EPR in other diseases with pathologically permeable vasculature.

## Introduction

Despite years of research and improvements in treatment strategies, malignant neoplasms remain the leading cause of death in adults ages 45-64 and the second leading cause of death for all age groups.^1^ This is true regardless of cancer research being one of the most well-funded disease topics through the NIH.^2^ There have been promising results in preclinical research using nanotherapeutics, but their translation from preclinical research and clinical trials to FDA approval has not been as successful as anticipated, with only 6 cancer nanotherapies approved as of 2019.^3^

The major benefits of nanomedicines are that they have reduced systemic side effects compared to free drug, and, in some patients, they can result in significantly greater drug deposition within the tumor and improved patient survival.^4,5^ However, most of these benefits have been demonstrated in animal models, not humans.^5–7^ Nanomedicines have shown significant inter- and intra-patient heterogeneity of therapeutic efficacy, which may contribute to these challenges.^8–10^ Despite this, some nanomedicines have shown enhanced therapeutic efficacy with patients.^5,11–13^

A primary cause of intra- and inter-patient variability in nanomedicine success is thought to be related to the heterogeneity of the tumor microenvironment itself.^9,10,14^ Many nanomedicines rely primarily on passive accumulation in tumors through the enhanced permeability and retention, or EPR, effect.^15^ Due to this, tumor characteristics like interstitial fluid pressure, vascular permeability, matrix density, and vascular supply can have a significant influence on the success of nanoparticles entering and remaining in the tumor interstitium.^16–25^ All of these characteristics can affect the success of passive nanoparticle accumulation; however, it remains difficult to determine who may be a good candidate for nanotherapy.

Stratifying patients into potential positive and negative responders for nanomedicine therapy is a strategy that has received increasing attention over the past decade to improve translation to the clinic.^5,26–29^ If patients can be pre-selected for nanomedicine treatment, the success rate of the therapeutic agent may also increase. There are multiple criteria for patient pre-selection, which can include histology and imaging biomarkers.^5^ Typically, with biopsy, only a small tissue sample is taken for pathology. Thus, the entire tumor microenvironment and its heterogeneity are not captured, and the sample has been removed from its native environment. In comparison, imaging biomarkers are non-invasive and can capture the entire heterogeneous tumor environment in real-time.^5,27,28^ The ideal imaging modality for this application would have a high safety profile with high temporal and spatial resolution. It would be quick, simple, inexpensive, and capable of performing repeat imaging studies as needed. Examples of imaging modalities include, but are not limited to, magnetic resonance imaging (MRI), computed tomography (CT), positron emission tomography (PET), optical imaging, and ultrasound.

Some imaging modalities have shown success in predicting nanoparticle extravasation based on tumor biomarkers and companion nanoparticles. Tumor biomarkers include vessel density, perfusion, angiogenic status, interstitial fluid pressure, collagen content, and more.^20,22–25,30–36^ Ultrasound has been used to measure many of these markers, especially vascular-related parameters, but imaging modalities including CT, MRI, and PET have also been applied. While tumor biomarkers have been able to predict nanoparticle extravasation in some studies, they may not be the ideal metric. Typically, these markers only apply to specific tumor types and measure a single metric in each test. Nanoparticle extravasation depends on many parameters; therefore, the probability of successful delivery to and retention in the tumor is based on a combination of factors. This probability may not be accurately captured by exclusively using anatomically based imaging biomarkers. A better approach could be using a companion nanoparticle.

A companion nanoparticle should be capable of being visualized through medical imaging and is used to predict whether a nanotherapeutic particle will be taken up and retained by the tumor.^37–40^ A successful companion nanoparticle would not impact the physiology contributing to nanoparticle uptake, and it would be cleared from the body quickly and safely. Companion nanoparticles should be designed to consider most of the barriers that therapeutic nanoparticles may encounter, making it potentially applicable to multiple tumor types and nanoparticle formulations. There have been numerous successes of companion nanoparticle strategies, particularly with MRI, PET, and CT.^11,24,26,33,37–46^ However, in realistic translation to the clinic, these imaging modalities have some disadvantages. CT and PET expose patients to ionizing radiation, which can be carcinogenic in large doses, adding a barrier to the ability to perform repeat studies.^47,48^ MRI is safe, but it has low temporal resolution, requires long scan times, and can be expensive.^49–52^ An imaging modality that hasn’t been used with companion nanoparticles yet is ultrasound. This modality is safe, inexpensive, quick, portable, capable of performing repeat studies, and has high temporal and spatial resolution.^53^ In analyzing the heterogeneity of tumors, high spatial resolution is important to identify regional differences in nanoparticle uptake. This is to determine if areas within the tumor may not be receptive to nanoparticles, in which case, alternative or combination strategies may be considered. Ultrasound can have a spatial resolution of about 200 microns for higher frequency scanners, which is superior to many other imaging modalities.^53^

Ultrasound image quality and soft tissue contrast can be improved by using contrast agents. Ultrasound contrast agents (UCAs) are bubbles in the nano- and micro-scale with shells made of lipids, polymers, or proteins.^53^ Microbubbles (MBs) are clinically approved blood pool agents, primarily used for vascular imaging.^53^ Nanoscale agents, or nanobubbles (NBs, ∼100-600 nm diameter), have recently emerged as an alternative to MBs (1-10 µm diameter).^53^ NBs are particularly well-suited to the role of a companion nanoparticle because they have been shown to extravasate out of tumor vasculature and into the tumor interstitium (assessed by histology and intravital microscopy).^54–59^

This work uses NBs as a contrast-enhanced ultrasound (CEUS)-based companion nanoparticle to differentiate human glioblastoma (U87) and colorectal adenocarcinoma (LS174T) tumors that have considerably different probabilities for nanoparticle extravasation due to their microenvironment characteristics.^60–64^ U87 tumors have small endothelial pores with poor nanoparticle penetration, while LS174T tumors have larger endothelial pores and higher nanoparticle penetration.^61,62,65^ Beyond differentiating these tumors with imaging biomarkers, this work also uses NBs to predict therapeutic doxorubicin-loaded liposome extravasation and retention and their heterogeneous deposition in tumors. Successful prediction of therapeutic nanoparticle extravasation and retention with NBs could improve the translation of nanoparticles from preclinical trials through clinical approval, thus improving patient care by pre-selecting patients who may be positive responders to cancer nanotherapy.

## Methods and Materials

### Nanobubble Production

NBs were fabricated according to published protocols (de Leon et al).^66^ Briefly, the lipids DPPE (1,2-dipalmitoyl-sn-glycero-3-phospho-ethanolamine; Corden Pharma, Liestal, Switzerland), DPPA (1,2-dipalmitoyl-sn-glycero-3-phosphate; Corden Pharma, Liestal, Switzerland), DSPE-mPEG-2k (1,2-distearoyl-snglycero-3-phosphoethanolamine-N-[methoxy(polyethylene glycol)-2000]; Laysan Lipids, Arab, AL), and DBPC (1,2-dibehenoyl-sn-glycero-3-phosphocholine; Avanti Polar Lipids Inc.; Pelham, AL)) were dissolved in propylene glycol (Sigma Aldrich; Milwaukee, WI) at 80°C via hot water bath and sonicated in an ultrasonic bath as needed (dependent on lipid dissolution rate). A solution of glycerol (Acros Organics) and phosphate-buffered saline (PBS; Gibco, Life Technologies) was simultaneously heated at 80°C via a hot water bath. The PBS-glycerol solution was added to the lipid solution 1 mL at a time while heated at 80°C. 1 mL of the mixed solution was added to a 3 mL headspace vial and sealed with a rubber stopper and aluminum cap. The headspace air was purged with 10 mL of C_3_F_8_ gas (AirGas; Cleveland, OH). NBs were activated via mechanical agitation using a VialMix mechanical shaker (Bristol-Meyers Squibb Medical Imaging Inc.; Billerica, MA) and then isolated via centrifugation at 50 g for 5 min. NBs were extracted from the centrifuged vial directly before use. NB concentration was approximately 4.07 x 10^11^ NBs/mL and had a mean diameter of 275 ± 8 nm.^66^ Measurements were obtained via resonance mass measurement (RMM; Archimedes, Malvern Panalytical Inc., Westborough, MA) and dynamic light scattering (DLS) (**Supplemental Fig. 1**).

### Cell Culture and Inoculation

U87 (human glioblastoma) cells were grown at 37°C and 5% CO_2_. They were maintained in complete Dulbecco’s Modified Eagle Medium (DMEM) media (Invitrogen Life Technology, Grand Island, NY). Male athymic nude mice (homozygous Foxn1^nu^) were anesthetized through inhalation of 2% isoflurane with 1.5 L/min oxygen and subcutaneously implanted (right hind limb) with 1 x 10^7^ U87 cells in 200 µL of Matrigel. LS174T (human colorectal adenocarcinoma) cells were grown at 37°C and 5% CO_2_ and maintained in complete Eagle’s Minimum Essential Medium (ATCC #30-2003). Male athymic nude mice were anesthetized through inhalation of 2% isoflurane with 1.5 L/min oxygen and were subcutaneously implanted (right hind limb) with 1 x 10^7^ LS174T cells in 30 µL of complete media (**Fig. 1A**). LS174T and U87 tumor types differ significantly in their vascular characteristics. LS174T tumors have a pore cutoff size of 400-600 nm while U87 tumors have a pore cutoff size of 7-100 nm, but their permeability to small molecules (e.g., albumin) is independent of pore cutoff size (**Fig. 1B**).^60,61^

**Fig. 1.**
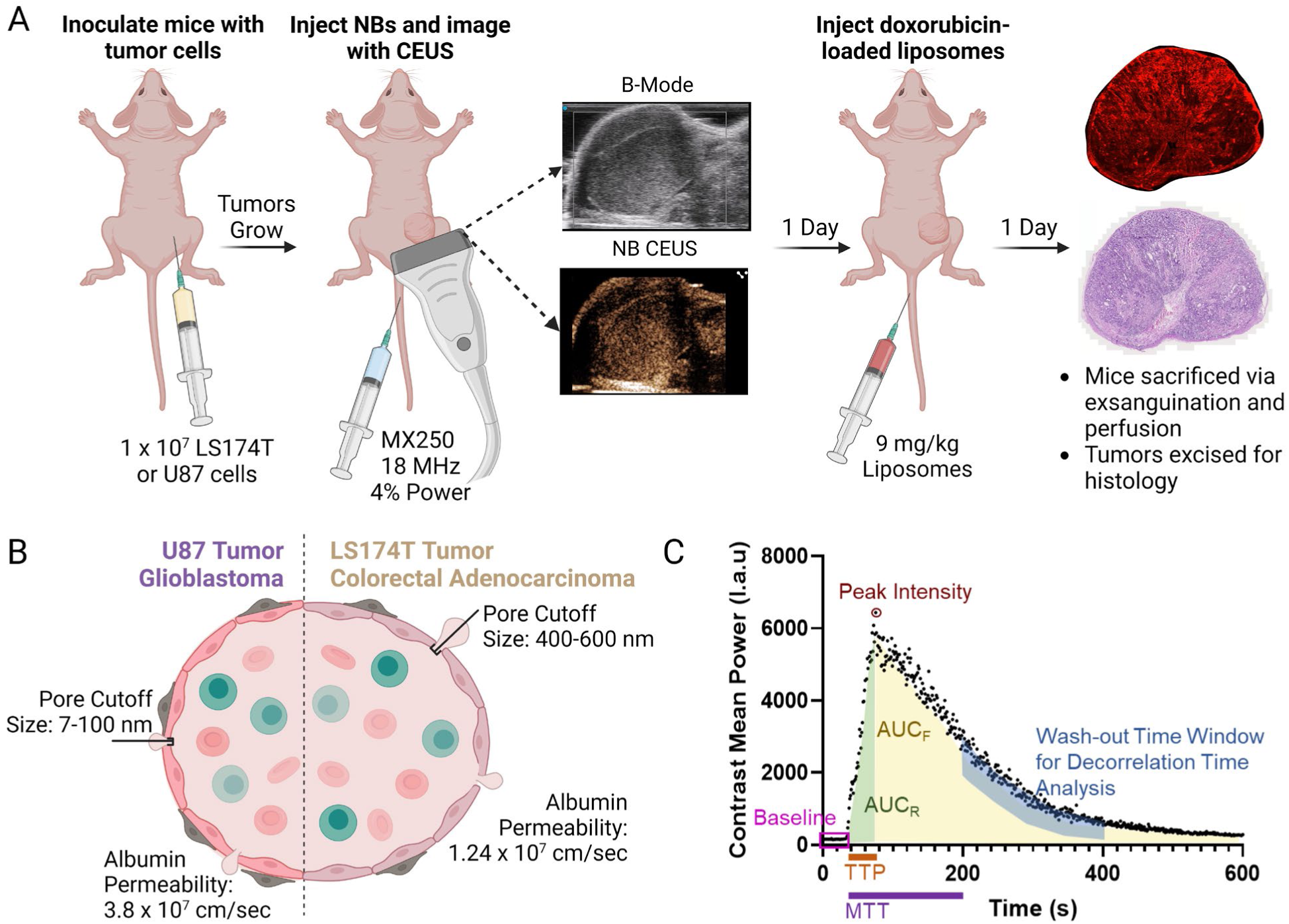
Experimental setup and parameters. (A) Mice were inoculated with tumor cells, and the tumors were permitted to grow until they reached at least approximately 250 mm^3^. Mice were then imaged with NB-contrast-enhanced ultrasound (CEUS) and B-mode imaging. 1 day post CEUS imaging, mice were injected with doxorubicin-loaded liposomes, which circulated for 24 hours before sacrifice via exsanguination. (B) Depiction of vascular differences between U87 and LS174T tumor types.^60^ (C) Graphical depiction of TIC parameters and their meaning based on Gu et al.^67^ DT analysis is based on work by Cooley et al. and Wegierak et al.^68–70^

15 LS174T mice and 15 U87 mice were inoculated for each cell line, but only 13 LS174T and 9 U87 mice were included in the analysis for reasons including no tumor growth (n = 1 U87), death under anesthesia (n = 1 LS174T), unsuccessful tail vein injection (n = 1 LS174T, n = 2 U87) and challenges related to imaging such as significant artifacts or loss of data before it was saved (n = 3 U87). Some tumor image analysis points were excluded due to the loss of wash-in data points, attenuation, and imaging artifacts. All tumor data used in this study can be found in **Supplemental Table 1**. A representation of the experimental flow can be seen in **Fig. 1A**.

### Animal Model

Male athymic nude mice were handled according to a protocol approved by the Institutional Animal Care and Use Committee (IACUC) at Case Western Reserve University and were in accordance with all applicable protocols and guidelines regarding animal use. Animals were observed twice a week until study termination. During imaging procedures, mice were under anesthesia through inhalation of 2% isoflurane with 1.5 L/min of air.

A subset of three mice was imaged weekly at Weeks 1 and 2 post-inoculation for LS174T tumors and at Weeks 2 and 3 post-inoculation for U87 tumors. The difference in the week was due to a slower tumor growth rate of U87 tumors. The mice were euthanized via exsanguination after the second week of imaging. For all other mice, imaging occurred once tumors reached a volume of at least approximately 250 mm^3^, although some tumors were smaller (**Supplemental Fig. 2**).

24 h after the final imaging time point, mice were injected via tail vein with 9 mg/kg of doxorubicin-loaded liposomes (Avanti Polar Lipids; Birmingham, AL) based on doxorubicin content (2.0 mg/mL), reported by the manufacturer. The excitation/emission spectrum of the liposomes (which guided the fluorescent imaging) can be found in **Supplemental Fig. 3**. The reported particle size, measured using DLS, was 95.5 nm or 109.3 nm, depending on the batch (300112-01-011 or 300112S-1EA-012). Liposomes were stored at -20°C until use. Doxorubicin-loaded liposomes were chosen to represent a model therapeutic nanoparticle both because they are similar to the commercially available Doxil® and because doxorubicin has natural autofluorescence.

Mice were euthanized 24 h after doxorubicin-loaded liposome injection via exsanguination to limit the number of liposomes within the tumor vasculature. During sacrifice, mice were anesthetized through inhalation of 2% isoflurane with 1.5 L/min oxygen. The exsanguination procedure involved puncture of the right atrium and perfusion of 50 mL of PBS through the left ventricle until the liver turned pale. The tumors were excised and stored in 4% paraformaldehyde for 24 hours. The tumors were then transferred to a 30% sucrose solution for 48 hours. They were embedded in OCT compound and stored at -80°C until sectioning. It should be noted that the tumors were embedded in OCT compound with labels corresponding to their *in vivo* orientation to make sure that tumor slice orientation corresponded as closely as possible to the ultrasound imaging plane and time-intensity curve (TIC) data.

### Ultrasound Parameters

A Vevo 3100 preclinical ultrasound imaging system (FUJIFILM VisualSonics; Toronto, ON) was used with an MX250 transducer (15-30 MHz bandwidth frequency) for all imaging studies. TIC images were acquired in dual B-mode and 2D nonlinear contrast (NLC) mode with an NLC transmit frequency of 18 MHz (4% power, 30 dB contrast gain, 28 dB gain, 40 dB dynamic range, 1 fps).^66,71^ The focus was aligned to the bottom of the tumor. All tumors were imaged with 3D imaging to measure tumor volume with Vevo LAB software (**Supplemental Fig. 2**). The ultrasound probe was attached to a 3D motor that was positioned as close to the center of the tumor as possible. Imaging commenced immediately after NB injection (200 µL via tail vein at approximately 300 µL/min) and continued for 30 minutes in one vertical plane of the tumor, still at the tumor center. Area was measured in the video acquisition plane and calculated by taking an ROI of the tumor in Vevo LAB software. All tumors were imaged with Doppler imaging in the same imaging plane as the TIC to identify major blood vessels.

### Time-Intensity Curve and Decorrelation Time Analysis

Regions of interest (ROIs) around the tumor were drawn using Vevo LAB software, which produced TICs of average signal intensity versus time. Tumor boundaries were chosen based on B-mode imaging and applied to NLC-mode videos. Some videos had areas with artifacts, seen with both B-mode and NLC imaging. Artifacts were excluded from TIC analysis. TIC analysis was performed using GraphPad Prism analysis software. TIC analysis included the following parameters: peak intensity (PI), time to peak intensity (TTP), area under the curve (AUC), area under the rising curve (AUC_R_), area under the falling curve (AUC_F_), mean transit time (MTT), and baseline intensity. Parameters were chosen based on those proposed by Gu et al.^67^ Baseline intensity was subtracted from all other TIC values for normalization. Imaging analysis began 1 second before NB entrance into the tumor.

Decorrelation time (DT) mapping was performed according to Wegierak et al.^69^ Briefly, greyscale videos of NLC mode imaging of the tumors were analyzed by pixel using MATLAB R2022a. DT mapping began at 50% of the peak intensity in the washout of the TIC with the intent of imaging NBs stuck in the vasculature or extravasated in the interstitium. Videos were analyzed for 200 frames, corresponding to 200 s. The correlation coefficient was thresholded at 0.5 based on prior studies.^68,70,72^ To find the average DT for the tumor, an ROI was drawn around the tumor in the DT map, and the pixel DTs were averaged using MATLAB R2022a.

### Histological Analysis

Each tumor was sectioned in 10 µm slices with samples from five evenly spaced sections of the tumor, including the center. Tissue samples were stained with H&E or were unstained (sectioning and staining through the Case Western Reserve University Tissue Resources Core). Tumors were sectioned in the same orientation as the plane of ultrasound imaging for comparison with TIC imaging. Unstained tumors were imaged using an Axioscan.Z1 (Zeiss Inc., Oberkochen, Germany) at 20x objective with 535 ms exposure time for each slide. Excitation wavelength filters were 456–483 nm and 543–568 nm. Emission wavelength filters were 497–527 nm and 581–679 nm. Note that not all tumors have corresponding histological data due to challenges with tail vein injection of the doxorubicin-loaded liposomes.

### Decorrelation Time Mapping Correlation with Histology

DT maps were compared to histological images using the corresponding quadrant. Histological sections were chosen to be approximately the same slice in the tumor as ultrasound imaging. This was done by comparing 3D ultrasound imaging to the five sections in the tumor where histological slices were taken. The histological slice most similar to the TIC imaging plane, by area, orientation, and morphology was chosen. This histological slice was most frequently one of the slides from the center of the tumor. Tumor quadrants were approximately even. Tumor area, calculated with Vevo LAB software for imaging data and ZEISS ZEN software for histological data, were also compared (**Supplemental Fig. 4**) to account for possible limitations in direct comparisons between imaging and histological analysis. Mean intensity values were measured for each section of the histological quadrant through ZEISS ZEN software. Tumors were excluded from analysis if their average DT was < 1 s due to temporal resolution limits caused by a 1 fps ultrasound imaging framerate.

### Statistical Analysis

TIC data was plotted and analyzed using GraphPad Prism. Unpaired parametric t-tests were used to directly compare LS174T and U87 tumor results. Linear regression was used to analyze fluorescent intensity versus average DT in each tumor quadrant. Analyses with a *p*-value of 0.05 were considered statistically significant.

## Results

### Time Intensity Curve Analysis

TICs, acquired through Vevo LAB software and analyzed using GraphPad Prism, were compared between LS174T and U87 tumors to determine if differences in TIC parameters could be ascertained through CEUS imaging. TICs were analyzed as an average intensity over the entire tumor ROI. Thus, they do not account for the intra-tumoral heterogeneity that is prominent in some tumors. Representative LS174T (**Fig. 2A-B**) and U87 (**Fig. 2C-D**) images in NLC and B-mode and corresponding TICs can be seen in **Fig. 2**. LS174T tumors showed higher average values for all parameters (**Fig. 3**). **Table 1** indicates the corresponding mean, standard deviation, and *p*-value for each parameter.

**Fig. 2.**
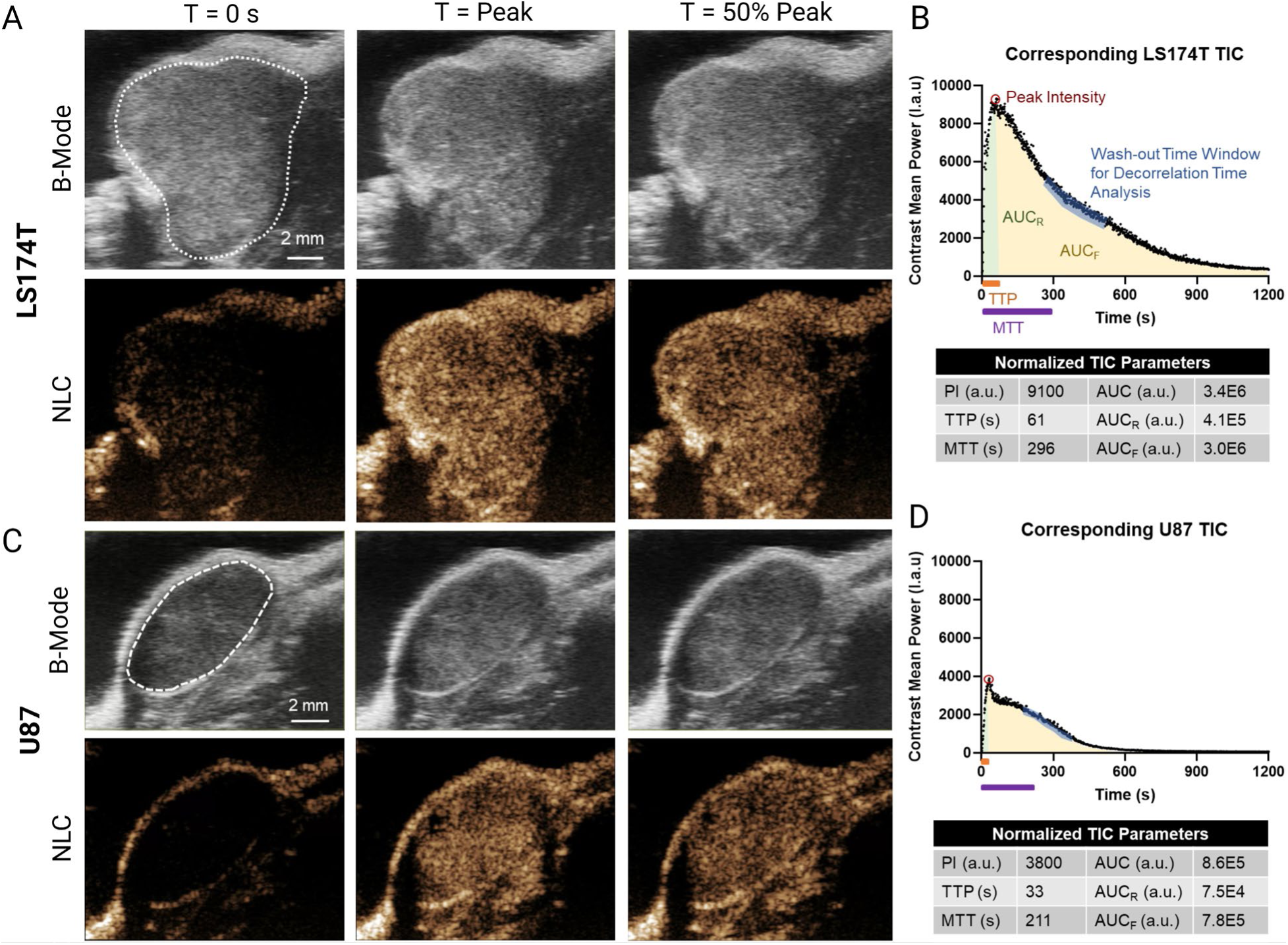
Representative TICs and Ultrasound Images of LS174T and U87 Tumors. (A) Representative B-mode and NLC mode images of an LS174T tumor at baseline (t = 0 s), peak intensity, and 50% of peak intensity in the wash-out. (B) Corresponding LS174T TIC and TIC-derived parameters, normalized by subtracting baseline intensity. (C) Representative B-mode and NLC mode images of a U87 tumor at baseline (t = 0 s), peak intensity, and 50% of peak intensity in the wash-out. (D) Corresponding U87 TIC and TIC-derived parameters, normalized to baseline intensity.

**Fig. 3.**
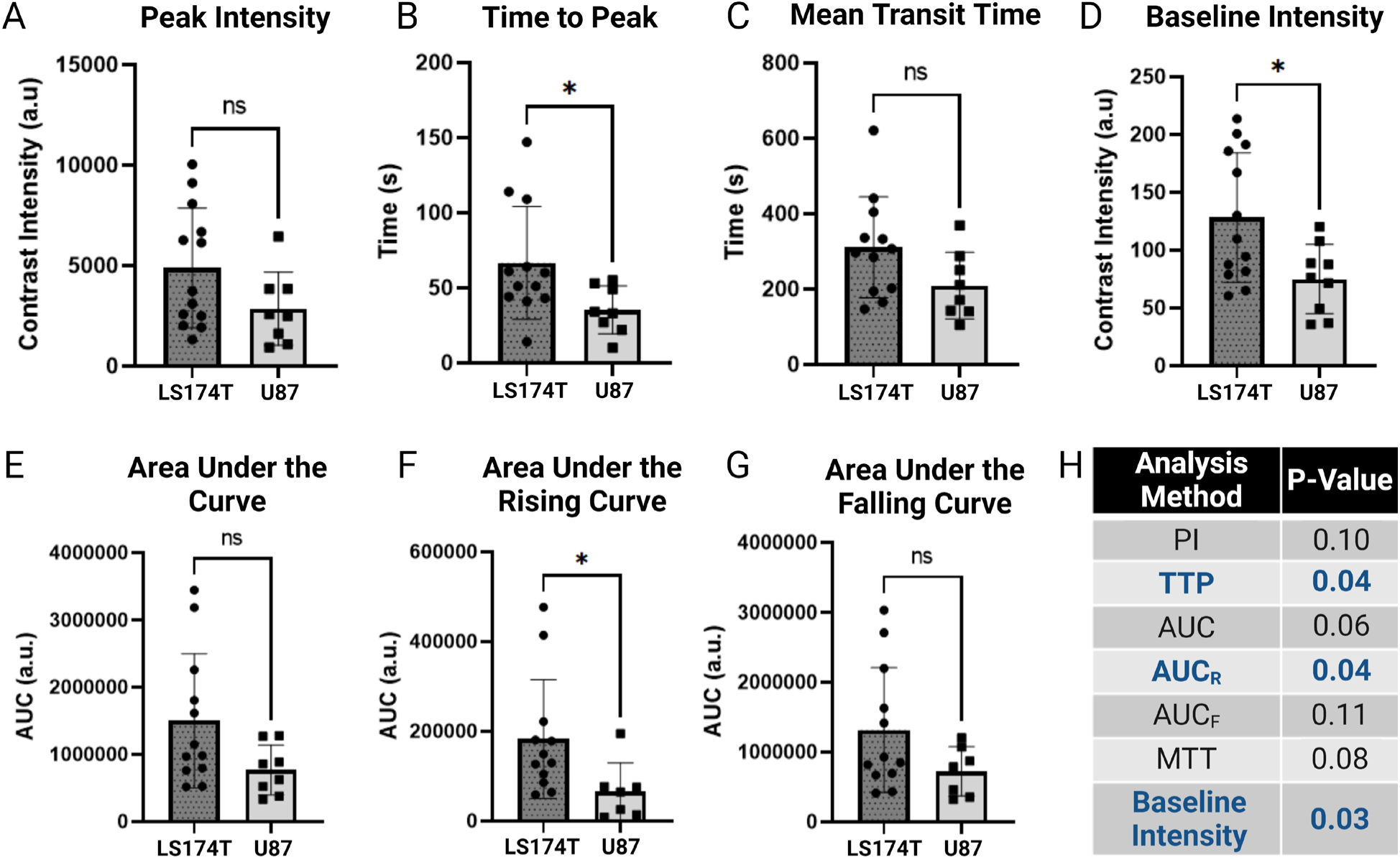
TIC Parameter Analysis between LS174T and U87 Tumors. (A) Peak intensity (PI). (B) Time to peak intensity (TTP). (C) Mean transit time (MTT). (D) Baseline intensity, or NLC intensity prior to NB entrance into the tumor. (E) Area under the curve (AUC). (F) Area under the rising curve (AUC_R_). (G) Area under the falling curve (AUC_F_). (H) P-values for all analyzed parameters. * indicates *p* < 0.05, and error bars represent standard deviation.

**Table 1.**
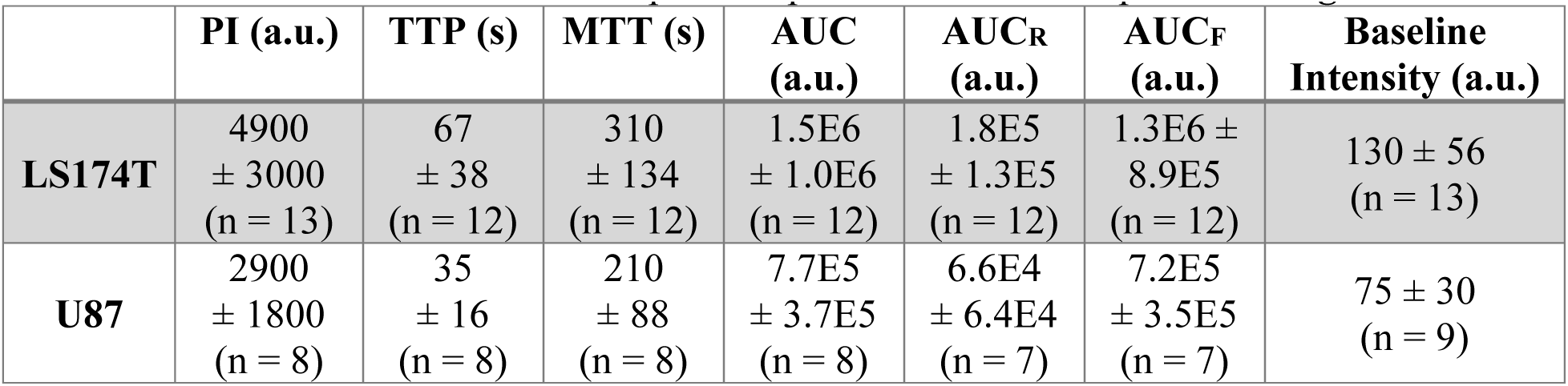
LS174T vs U87 TIC comparison values. LS174T and U87 mean and standard deviation values are indicated as well as their comparative p-values. Values are plotted in **Fig. 3**.

Of note, TTP, AUC_R_, and baseline intensity showed statistically significant differences between tumor types. On average, TTP was 1.9x greater in LS174T tumors than U87 tumors, AUC_R_ was 2.7x greater, and baseline intensity was 1.7x greater. Despite not being statistically significant, all other parameters showed a higher mean value for LS174T tumors compared to U87 tumors. PI was 1.7x greater, MTT was 1.5x greater, AUC was 1.9x greater, and AUC_F_ was 1.8x greater. All tested parameters had large standard deviations, with LS174T tumors showing more variation than U87 tumors. Predictably, tumors were highly heterogeneous between mice and within the same tumor, so large standard deviations were expected.

### Decorrelation Time Analysis

DT was analyzed at 50% of the PI in the wash-out of the TIC for each tumor. This time point was chosen to represent patterns in NB movement after NBs were given sufficient time to enter pathological blood vessels and begin extravasation. DT maps were generated through MATLAB, and ROI pixel values were averaged. When LS174T and U87 tumors were compared by averaging all pixels within the tumor, the LS174T tumors had a statistically (*p* = 0.04) greater average DT (**Fig. 4A**). LS174T tumors (n = 12) had an average DT of 2.6 ± 1.2 s while U87 tumors (n = 8) had an average of 1.4 ± 1.2 s. It should be noted that an outlier of 11.0 s was excluded from the LS174T group. When the standard deviation of the DT of each intratumoral pixel DT was compared, LS174T tumors (n = 12) had a significantly greater (*p* = 0.04) standard deviation compared to U87 tumors (n = 8) (**Fig. 4B**). This indicates that the LS174T tumors had a more variable DT within the tumor, suggesting higher intratumoral heterogeneity.

**Fig. 4.**
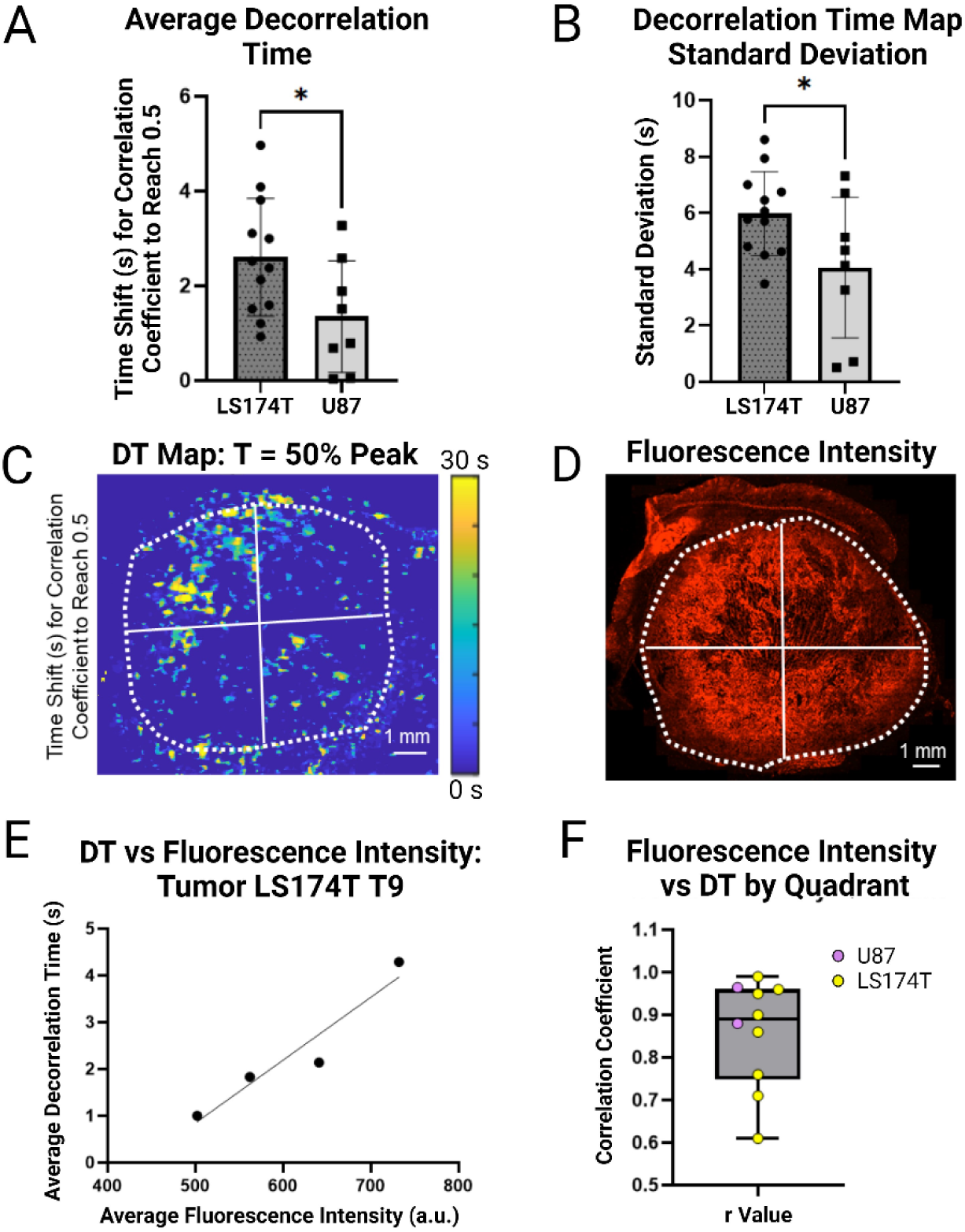
Decorrelation Time (DT) Analysis. (A) DT for each tumor found by averaging the DT map pixels within the tumor ROI. (B) Standard deviation of DT between all intratumoral pixels. (C) Representative DT map split into quadrants. (D) Doxorubicin-generated fluorescent intensity map from the same tumor and approximately the same imaging plane as Fig. 4C. The tumor is divided into corresponding quadrants. (E) Representation of the linear regression of the average DT vs average fluorescence intensity, corresponding to the tumor in Fig. 4C-D. (F) Correlation coefficient (r) of all tumors with an average DT (Fig. 4A) > 1 s that had corresponding histological data available. Corresponding quadrants of DT and fluorescence intensity, as in Fig. 4C-D, were compared to generate the correlation coefficients. U87 and LS174T tumors are labeled. *indicates *p* < 0.05 and error bars represent standard deviation.

### Decorrelation Time vs Fluorescent Intensity of Doxorubicin

Due to the high intratumoral variability of DT, tumors were split into quadrants and directly compared to fluorescence intensity in corresponding quadrants on histology. Quadrant-based comparison allowed us to determine if the heterogeneity in DT maps resulted from tumor physiology or was a methodological artifact. This quadrant analysis method used each histological slide as its own control, reducing the effect of the background signal between slides. A representative tumor can be seen in **Fig. 4C-D**, with the corresponding quadrants drawn. The quadrants were compared through linear regression, generating a correlation coefficient. The representative tumor in **Fig. 4E** had a correlation coefficient of r = 0.96. This analysis was performed for all tumors with an average DT > 1 s (n = 8 LS174T, n = 2 U87) (**Fig. 4A**). Tumors with an average DT < 1 s were excluded because the imaging frame rate was 1 fps. A higher frame rate is required to capture accurate data for tumors with low DTs. The average correlation coefficient for all included tumors was 0.86 ± 0.13 (**Fig. 4F**). It should be noted that the imaging plane and histological slice were never perfectly aligned, thus accounting for some variance. The area of the imaging plane and histological slice were compared for LS174T tumors. There was a negative correlation coefficient of -0.74 between r value and % difference in tumor area (**Supplemental Fig. 4**). Thus, as the percent difference in tumor area increased, the correlation coefficient also decreased, indicating that a mismatch of the imaging plane and histological slice could be why some correlation coefficients did not show as strong of a correlation as others.

In **Fig. 5**, an LS174T tumor is presented, further illustrating intratumoral heterogeneity. Different tumor regions may have substantially different TICs owing to differences in vascular permeability, density, and flow patterns (**Fig. 5A, D-F**). The heterogeneous tumor physiology is evident through histology with H&E staining and doxorubicin-liposome generated fluorescence intensity data (**Fig. 5B-C**). Between Regions 1 and 2, the average PI was 4.9x higher for Region 2, while the TTP was 7.6x greater for Region 1. Thus, while producing an average TIC from an entire tumor is a simple metric for intertumoral comparison, it does not always accurately capture how each tumor region will respond to drug-loaded nanoparticle injection. Quadrant analysis and mapping are more accurate ways to measure the heterogeneity of the tumor and relate it to nanoparticle distribution.

**Fig. 5.**
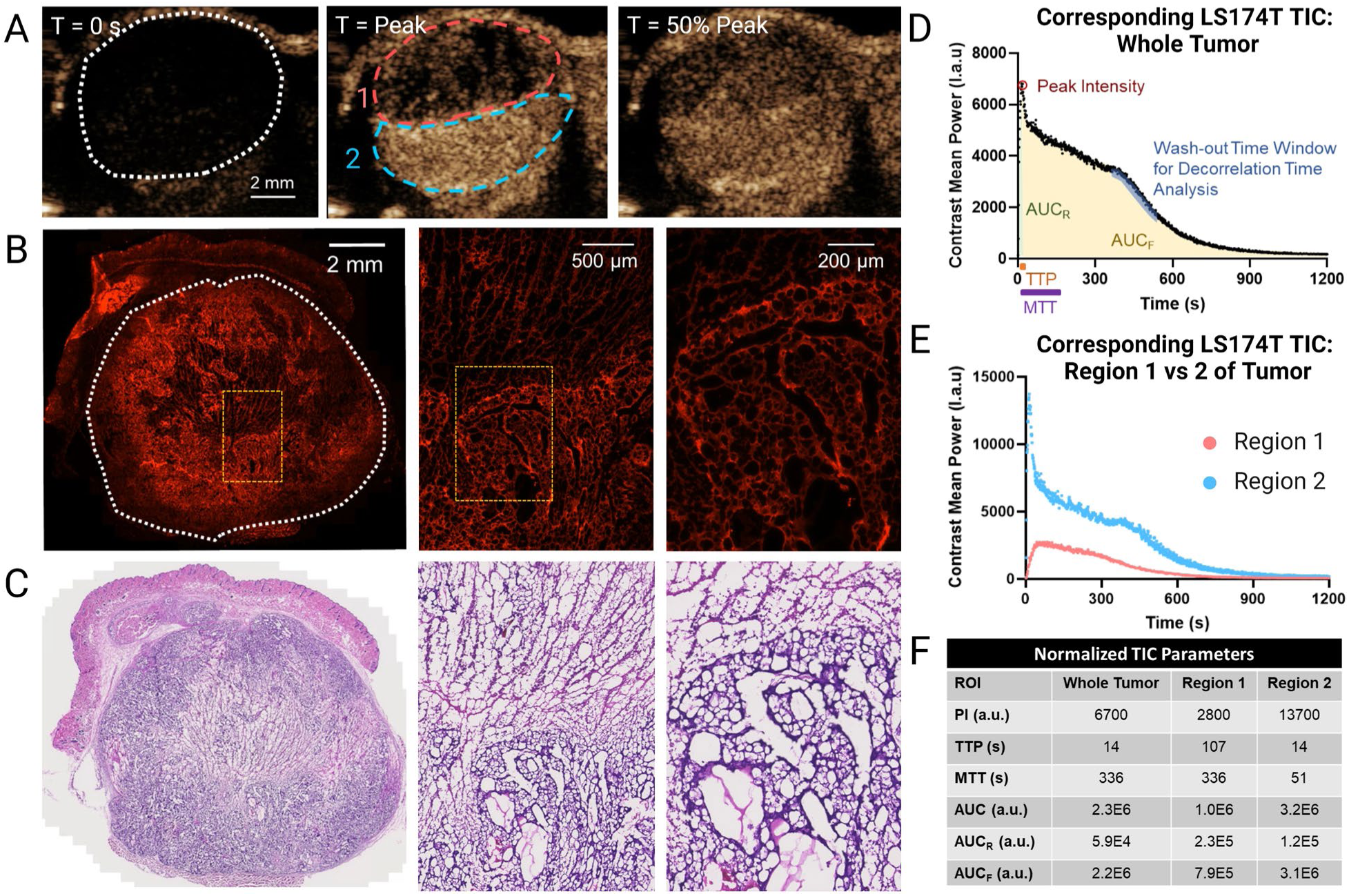
Representation of Intratumoral Heterogeneity. (A) NLC of an LS174T tumor at baseline (t = 0 s), peak intensity, and 50% of the peak intensity in the washout. The ROIs of the tumor have been split into the entire tumor (white dashed lines), Region 1 (pink dashed lines), and Region 2 (blue dashed lines). (B) Fluorescence intensity originating from doxorubicin-loaded liposomes of the corresponding tumor, zoomed in to the yellow ROI in each subsequent image. (C) H&E histological image of the corresponding tumor with the same magnification and ROI as Fig. 5B. (D) TIC of the ROI of the whole tumor (white dashed lines). (E) ROIs when the tumor is divided into Regions 1 and 2. (F) Normalized TIC parameters for each ROI.

### Relation of DT and TIC to Tumor Progression

As tumors grow, they must develop their own vasculature, which is often pathological. For tumors, this angiogenic process begins a few days after implantation and continues as the tumor grows and requires more nutrients.^73^ Three LS174T and U87 tumors were imaged for two consecutive weeks and TIC and DT maps were generated (**Fig. 6A-F**). The weekly differences are evident in the shape of the TIC curves for each tumor, with all LS174T tumors having a larger PI and AUC for the second week of imaging compared to the first. U87 tumors were more variable, with a mix of TIC parameters between the first and second week of imaging. The average DT of all LS174T and 2/3 U87 tumors increased from the first to the second week of imaging (**Fig. 6G-H**). The average slope for LS174T tumors between Weeks 1 and 2 of imaging was 2.5 ± 1.6 (s/week), and the average slope for U87 tumors between Weeks 2 and 3 was 0.6 ± 0.7 (s/week).

**Fig. 6.**
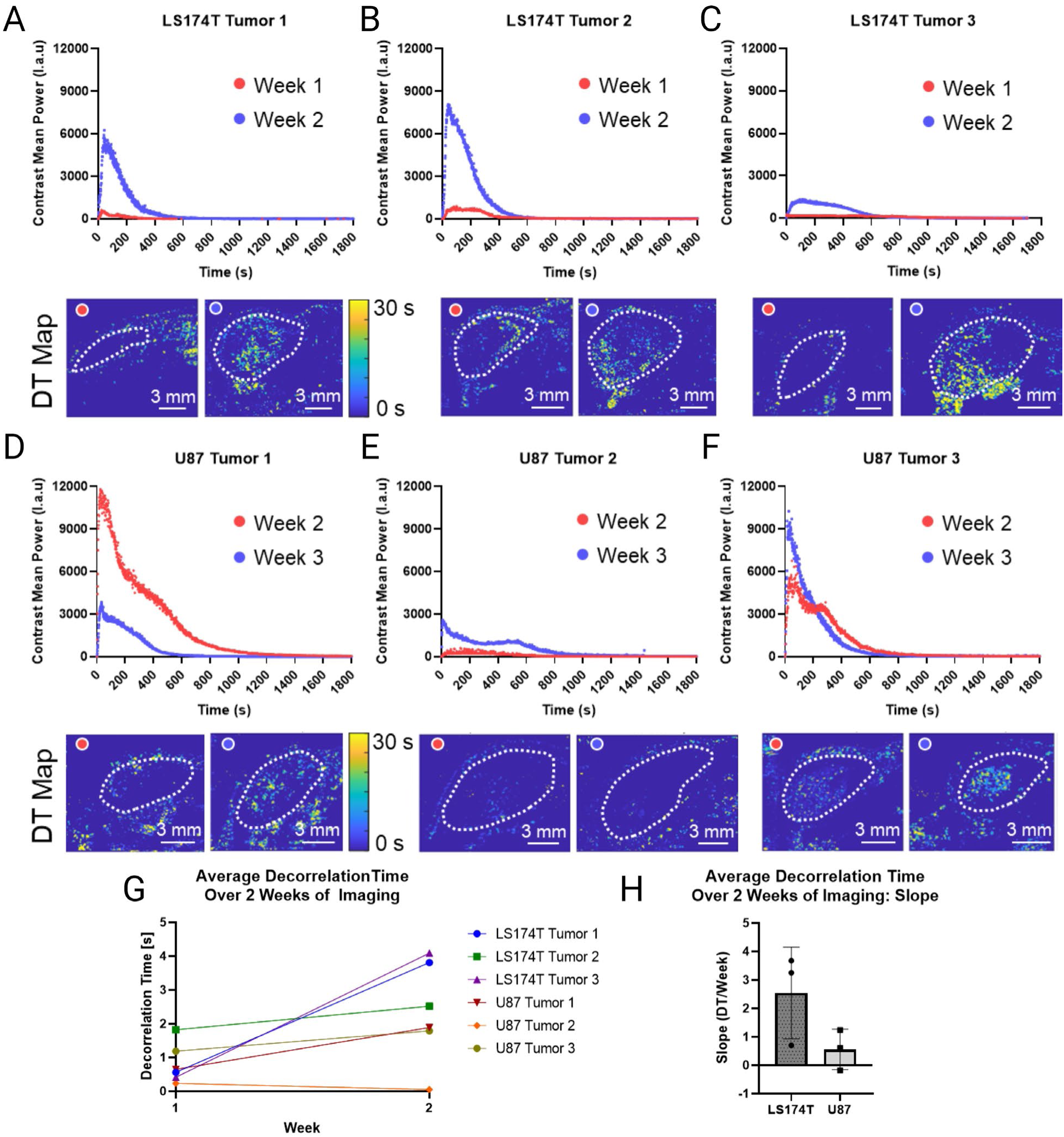
TICs and DT Maps of LS174T and U87 Tumors over 2 Weeks of Growth. (A-C) TICs and DT maps of three LS174T tumors between Week 1 and Week 2 of growth post-inoculation. (D-F) TICs and DT maps of three U87 tumors between Week 2 and Week 3 of growth post-inoculation. (G) Average DT of each tumor between the first and second week of imaging. (H) Slope of average DT between the first and second week of imaging.

## Discussion

Tumor heterogeneity remains a major challenge for nanomedicine-based therapeutics.^4,10,13^ This issue can be, in part, addressed through patient stratification strategies using imaging biomarkers with companion nanoparticles.^5,28^ Imaging biomarkers are non-invasive and can capture the morphology and physiology of the entire heterogeneous tumor in its native environment, potentially making it the preferred strategy. Companion nanoparticles have previously been used with MRI, PET, and CT, but haven’t been proposed using ultrasound until now.^28,37,39,40^ Nanobubble-based dynamic contrast-enhanced ultrasound, or NB-CEUS, is the ideal method for an ultrasound-based companion nanoparticle because NBs have been shown to extravasate from the tumor vasculature into the interstitium through intravital microscopy and histology.^54–56,58,74^ This study used NB-CEUS to differentiate two tumor types and related imaging biomarkers to doxorubicin-loaded therapeutic nanoparticle tumoral distribution through histology.

LS174T and U87 cell lines were chosen for their differing vascular characteristics. Based on a study by Hobbs et al., these tumor types differ significantly in their pore cutoff sizes, a measure of transvascular gap size, with LS174T tumors (400-600 nm) having a much larger pore size than U87 tumors (7-100 nm).^60,61^ Furthermore, the study by Hobbs et al. examined albumin permeability in both tumor types and found no significant differences, implying that permeability to small molecules is independent of transvascular gap size.^60^ Thus, the major difference between the LS174T and U87 tumors is that larger particles are more likely to escape from the intravascular space of the LS174T tumors. Another study used confocal microscopy to analyze 32 tumor models, including U87 tumors and a colorectal adenocarcinoma cell line (HT-29) tumor.^62^ They found that U87 tumors had one of the lowest amounts of nanoparticle (12 nm ferritin nanocage) penetration into the tumor interstitium out of the 32 tumor types, despite having a relatively high vascular density. This highlights how vascular density does not necessarily correlate directly to permeability. In contrast, the colorectal adenocarcinoma had the highest amount of nanoparticle penetration.^62^ A benefit to choosing LS174T cells as opposed to a cell line that results in tumors with greater permeability (such as an MCa IV murine mammary carcinoma with a pore cutoff size of 1.2-2 µm) is that the LS174T cell line is a more realistic study for a medium permeability tumor, which is more comparable to human tumors.^60^ Thus, LS174T and U87 tumors can be used to compare tumors with higher vs lower probability for nanoparticle extravasation.

To compare the contrast agent dynamics in LS174T and U87 tumors using ultrasound imaging, TIC parameters were calculated by averaging the NB generated signal intensity within an ROI of the tumor. Tumor boundaries were selected based on boundaries identified in B-mode imaging (**Fig. 2**). On average, LS174T tumors had higher mean values for *all* TIC parameters compared to U87 tumors. However, data spread for both tumor types was significant, likely due to the tumor heterogeneity, especially with the tumor vasculature, which is known to be highly spatially heterogeneous.^75,76^ CEUS imaging occurs in real time where the wash-in of the NBs through the vessels is visible. Thus, TIC parameters, which are dependent on NB kinetics within the tumor, also depend on tumor vascular location and integrity. In this study tumor TICs were acquired in only one imaging plane. Thus, the entire tumor, and its heterogeneity, may not be accurately represented. Furthermore, this analysis takes an average intensity value at each time point, averaged over the ROI. Hence, it does not capture the apparent spatial heterogeneity in some tumors (**Fig. 5**).

Wash-in parameters, including TTP and AUC_R_ (**Fig. 3B, F**), were significantly greater for LS174T tumors. Wash-in kinetics have been used to evaluate the extent of capillary leakage and differentiate tumor types with a non-nanoparticle dye as well.^77^ TTP measures how long it takes from the entrance of NBs into the tumor to when the tumoral contrast intensity reaches a peak. AUC_R_ takes into account both TTP and PI. Thus, vascular parameters like slow flow, tortuosity, blind ends, and functional blood vessel density may directly affect both TTP and AUC_R_.^78–80^ If a tumor has tortuous or leaky vessels, it may take longer for NBs to enter and exit the tumor. This increases the TTP and the potential total number of NBs that can enter the tumor, increasing PI and AUC_R_. Importantly, TTP and AUC_R_ are simple parameters that are easy to compute, potentially making them easy to apply to clinical practice.

MTT is another parameter likely affected by the vascular integrity of the tumor and vascular morphology (**Fig. 3C**). The average MTT was 1.5x greater for LS174T tumors compared to U87 tumors, although this difference was not statistically significant. However, with MTT, unlike with TTP, the continuous insonation under ultrasound could destroy extravasated NBs immobilized in the imaging plane. NBs that appear transiently in the imaging plane during normal vascular flow will not be as affected by acoustically-driven gas diffusion as NBs that are immobilized in the interstitium or poorly formed vessels continuously exposed to the ultrasound beam.^81^ NBs that have entered the interstitium or remain in slowly flowing vessels will eventually lose their echogenicity. Thus, if the NBs in the LS174T tumors are immobilized in a single location, they may dissipate more rapidly than NBs transiently exposed to ultrasound in the imaging plane.

PI is also highly dependent on tumor heterogeneity. **Fig. 5A, D-F** shows that the rate of reaching PI can vary by tumoral spatial location. In the tumor highlighted in **Fig. 5**, TTP is 7.6x longer in Region 1 while PI is 4.9x greater in Region 2. These results are likely due to regional differential NB access and accumulation within the tumor or a difference in maximum NB concentration. Hence, the average PI across the entire tumor does not reflect the spatial heterogeneity within the tumor. Importantly, **Fig. 5** shows that NB-CEUS can be used to identify and evaluate tumor heterogeneity. This knowledge could allow physicians to select successful treatment courses more accurately, including those that involve multiple therapeutic strategies (e.g., a nanoparticle in combination with a traditional chemotherapeutic or radiotherapy).

The AUC (**Fig. 3E**) is dependent on many factors, including residency time within the tumor and PI. It should be noted that individual NBs cannot be resolved by ultrasound with the acquisition parameters and hardware used in this study. Thus, small populations of NBs may not contribute to the AUC. It may be beneficial for a future study to target NBs to tumor cells in two tumor types that exhibit differing levels of permeability, similar to the setup of this study. The low permeability tumor would likely have a significantly lower AUC than the higher permeability tumor because, despite targeting, the NBs cannot reach the cells if they cannot escape the blood vessels. The AUC_F_ (**Fig. 3G**) makes up the largest proportion of the AUC (∼80-95% for most tumors), compared to AUC_R_ (**Fig. 3F**), so, understandably, this value has a similar trend to AUC.

The next parameter in this work was decorrelation time (DT), which was recently shown to be a novel NB CEUS-based biomarker specific to tumor detection.^69^ DT captures extravascular NB entrapment based on time-series CEUS data. In this analysis, LS174T tumors had a significantly longer average DT than U87 tumors (**Fig. 4A**). DT has been previously studied with NBs and found that when NBs are in restrictive environments (e.g., surrounded by blood cells in a static environment or diffusing through an extracellular matrix in a microfluidic model), these pixels have a longer DT.^68–70,82^ For NBs in blood vessels, DT will be close to zero if they move faster than can be captured by imaging data (limited by framerate). In this study, images are acquired at 1 fps, so the NB agent would need to stay in the imaging voxel for at least 1 second to produce a non-zero DT.^69^ Higher imaging frame rates could generate a greater range of DT data in future studies. Additionally, the ultrasound instrument limits the detail and resolution of DT maps. Thus, the accuracy of the DT maps may be hindered at lower-frequency imaging parameters.

While pixel analysis helps to account for some heterogeneity, another way to quantify this data is to divide the tumor into quadrants and analyze the average DT for each quadrant. Averaging is required for comparison to histology because of the differences in ultrasound and histological resolution. The therapeutic doxorubicin-loaded liposome was given 24 hours to circulate before sacrifice to ensure maximum accumulation, which is reported to reach a peak around 24 hours after injection.^83^ It should be noted that the autofluorescence signal of the tumor tissue was variable between tumors, which is another benefit of the internal control of performing a quadrant analysis. Autofluorescence signal on histological analysis has also been a challenge in other studies.^84^ Tumor sections for histology were taken at five evenly spaced sections along the tumor. These sections were matched as well as possible to imaging data by comparing each section to 3D ultrasound imaging (**Supplemental Fig. 4**).

There was a positive linear correlation between average DT and fluorescence intensity with an average correlation coefficient of 0.86 ± 0.13 (**Fig. 4F**). As noted above, the tumors with the highest correlation were also well matched in tumor area between ultrasound imaging and microscopy. An explanation for a lower correlation coefficient in some tumors is imperfect matching. This analysis implies that DT mapping can be used to predict liposomal nanoparticle EPR in a tumor after 24 hours of circulation. Thus, this study indicates that NBs can be a companion nanoparticle to predict therapeutic nanoparticle extravasation and retention. Future work should include TIC data with multiple simultaneously acquired imaging planes to make a 3D DT map. This is possible through quantitative 3D dynamic CEUS and could be used to identify if a whole tumor or regions of the tumor are susceptible to nanoparticle-based treatment.^85,86^ To this end, patients could be selected to receive nanoparticle therapy or they could receive nanoparticle therapy for one region of their tumor and another therapy (e.g., radiotherapy) in a different region.

This study also enabled a limited longitudinal analysis of TIC parameters with tumor progression. TICs changed significantly within 2 weeks, with the most notable difference in contrast dynamics seen in the DT parameter. DT increased to a greater extent with time for the LS174T tumors compared to the U87 tumors. In **Fig. 6**, average DT was measured between two weeks of growth for each tumor type and found that for 5/6 tumors, DT increased between the first and second week of imaging. This suggests that vascular permeability can increase with tumor progression, consistent with prior studies.^87^

The last parameter significantly different between U87 and LS174T tumors was baseline intensity (**Fig. 3D**) before contrast injection. While the mechanism for this is unclear, LS174T tumors exhibited a higher baseline contrast intensity than U87 tumors. Baseline NLC signal differences, and their mechanism, should be investigated in future work as potential biomarkers.

In addition to identifying when tumor vasculature becomes permeable, a future application of NB-CEUS could be to monitor treatment efficacy of therapeutic nanoparticles or traditional chemotherapeutics over time. Recommended treatment course may change as tumors grow or shrink. One group found that nanoparticle accumulation in tumors can become more heterogeneous over time as treatment progresses and other groups have also identified a need for repeat imaging to monitor treatment progress (e.g., tumor size, therapeutic efficacy).^42,44,88^ Ultrasound is ideal for repeat imaging due to its low cost and safety profile.

There are some limitations to this study. Imaging data was acquired in only one imaging plane; thus, the tumor heterogeneity throughout the entire volume was not captured. This study is the first proof of concept that NB-CEUS imaging data relates to tumor microenvironment characteristics and therapeutic nanoparticle distribution. However, future work could adapt a matrix array transducer for NLC imaging to capture 3D data in real-time.^89,90^ Additionally, pixel-based mapping of all TIC parameters could better account for heterogeneity than the averaged TICs presented in this paper. While nanoparticle access and accumulation are important for predicting treatment efficacy, future experiments will apply these ultrasound metrics to the efficacy of a therapeutic nanoparticle to predict tumor growth and animal survival. While this study focuses on the oncological applications of NB-CEUS, the strategies employed in this work could be applied to any pathology involving permeable vasculature, including cardiovascular diseases, the diabetic pancreas, sepsis, and more.

## Conclusion

This study reports the first ultrasound-based companion nanoparticle capable of providing information about tumor vascular permeability and the extravascular space. LS174T and U87 tumors were distinguished through multiple imaging parameters, including AUC_R_, TTP, DT, and baseline signal intensity. The NB-CEUS imaging biomarkers explored herein successfully distinguish between two tumor types and established NBs as a potential companion nanoparticle. When analyzed by tumor quadrant, LS174T and U87 tumors showed a strong positive correlation between DT and therapeutic nanoparticle distribution, suggesting that DT maps can accurately predict nanoparticle tumor accumulation. These findings could ultimately be applied to stratifying patient tumors into positive and negative responders to nanomedicine therapies for improved therapeutic efficacy. Due to the safety profile, versatility, low cost, and the proof of concept presented in this work that NBs are suitable companion nanoparticles, NB-CEUS may be the future of therapeutic nanoparticle-based personalized medicine in clinical oncology.

## Acknowledgements

This research was supported by the National Heart, Lung, and Blood Institute (F30HL160111); the National Institute of Biomedical Imaging and Bioengineering (R01EB025741 and R01EB028144); and the National Institute of General Medical Sciences (T32GM007250). F. Berg was supported by Marcos Lottenberg and Marcus Wolosker International Fellowship for Physicians Scientist. All figures were created using BioRender.com.

