## Supplemental Files for "Assessing Tumor Microenvironment Characteristics and Stratifying EPR with a Nanobubble Companion Nanoparticle via Contrast-Enhanced Ultrasound Imaging"

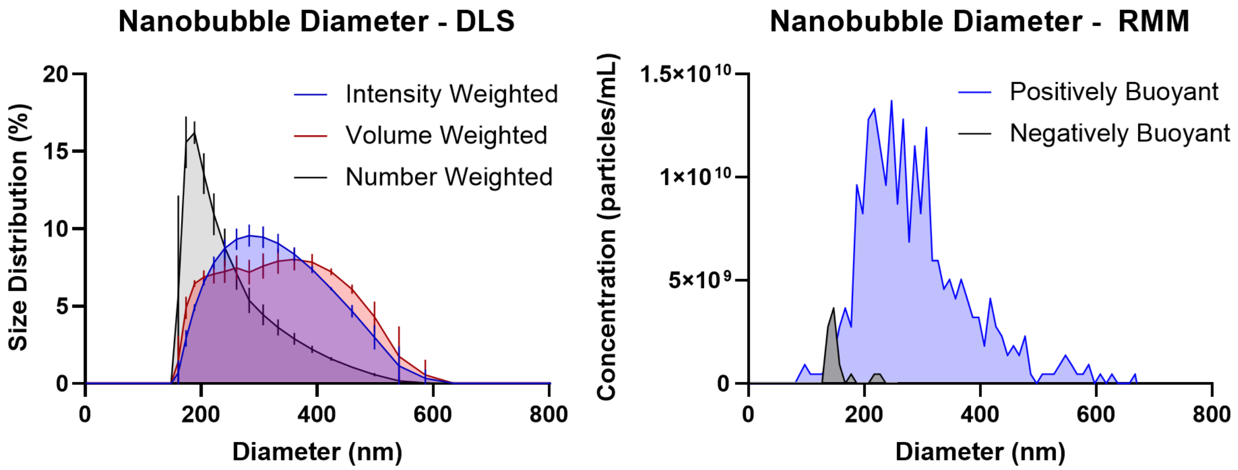


**Supplemental Fig. 1**. **Representative nanobubble diameter measurements using DLS and RMM.** Left: DLS measurement (n = 3) of nanobubble diameter through intensity, volume, and number weighted measurements. The average hydrodynamic radius was 306 ± 4. Right: RMM measurement (n = 1) of nanobubble diameter. The concentration of positively buoyant particles was 2.2 x $10^{11}$ nanobubbles/mL with a mean diameter of 290 ± 100 nm. The concentration of negatively buoyant particles was 8.7 x $10^{9}$ particles/mL with a mean diameter of 160 ± 30 nm.

| Tumor | Volume (mm^3) | Area (mm^2) | PI (a.u) | TTP (s) | AUC (a.u) | AUCr (a.u) | AUCf (a.u) | MTT (s) | Baseline Intensity (a.u) | DT 50% washout (s) | DT STD (s) |
|---|---|---|---|---|---|---|---|---|---|---|---|
| LS174T T1 | 662.455 | 45.968 | 6269 | 44 | 980998 | 129884 | 848750 | 147 | 167.193 | 3.8115 | 7.0074 |
| LS174T T2 | 1105.827 | 74.519 | 8076 | 43 | 1613948 | 180704 | 1416074 | 202 | 185.783 | 2.52 | 5.7701 |
| LS174T T3 | 469.469 | 31.171 | 6141 | 51 | 1806510 | 178281 | 1628229 | 307 | 81.787 | 1.59 | 4.8005 |
| LS174T T4 | 777.082 | 42.428 | 1321 | 114 | 524804 | 104746 | 410057 | 441 | 60.473 | 4.088 | 7.9397 |
| LS174T T5 | 978.175 | 76.392 | 2018 | X | X | X | X | X | 213.616 | 2.9999 | 6.7466 |
| LS174T T6 | 228.375 | 26.507 | 3734 | 64 | 793955 | 126515 | 667440 | 194 | 94.502 | 3.1058 | 6.4583 |
| LS174T T7 | 230.124 | 23.093 | 3108 | 51 | 518485 | 85978 | 432507 | 165 | 109.662 | 10.9536 | 11.5262 |
| LS174T T8 | 585.802 | 71.537 | 1934 | 147 | 1148580 | 221805 | 926775 | 621 | 65.044 | 0.9252 | 3.4892 |
| LS174T T9 | 948.292 | 76.495 | 6680 | 14 | 2257849 | 58851 | 2198997 | 336 | 130.115 | 2.38 | 6.06 |
| LS174T T10 | 1139.83 | 82.365 | 2496 | 41 | 760321 | 64431 | 695893 | 333 | 87.617 | 4.97 | 8.6 |
| LS174T T11 | 534 | 46.809 | 10048 | 60 | 3183522 | 476532 | 2706990 | 285 | 191.271 | 1.51 | 4.63 |
| LS174T T12 | 451.114 | 47.762 | 2579 | 109 | 968139 | 149383 | 818756 | 404 | 78.875 | 2.13 | 5.7 |
| LS174T T13 | 811.448 | 72.443 | 9122 | 61 | 3444956 | 414246 | 3030710 | 296 | 200.701 | 1.2 | 4.5 |
| U87 T1 | 60.637 | 8.766 | 6446 | 55 | 1267142 | 194843 | 1072299 | 171 | 87.634 | 3.2727 | 7.3084 |
| U87 T2 | 42.493 | 9.405 | 928.5 | 22 | 331353 | 8794 | 322558 | 250 | 76.341 | 1.512 | 4.6805 |
| U87 T3 | 328.998 | 31.577 | 3844 | 33 | 859134 | 75455 | 783676 | 211 | 49.517 | 1.8861 | 5.139 |
| U87 T4 | 557 | 51.704 | 2587 | 10 | 886241 | 14131 | 872110 | 141 | 36.995 | 0.0509 | 0.7212 |
| U87 T5 | 89.431 | 15.308 | X | 27 | X | X | X | 105 | 107.846 | 0.7895 | 3.2639 |
| U87 T6 | 359.749 | 23.505 | 1623 | X | 626126 | X | X | X | 35.749 | X | X |
| U87 T7 | 255.215 | 16.642 | 3846 | 34 | 1277369 | 76882 | 1200487 | 287 | 120.259 | 0.0331 | 0.5085 |
| U87 T8 | 1922.66 | 114.6 | 1090 | 49 | 377774 | 25566 | 352209 | 369 | 88.662 | 2.58 | 6.7065 |
| U87 T9 | 1320.334 | 113.919 | 2494 | 53 | 520016 | 64755 | 457637 | 143 | 71.621 | 0.6833 | 4.1345 |

**Supplemental Table 1**. All tumor analysis data points.

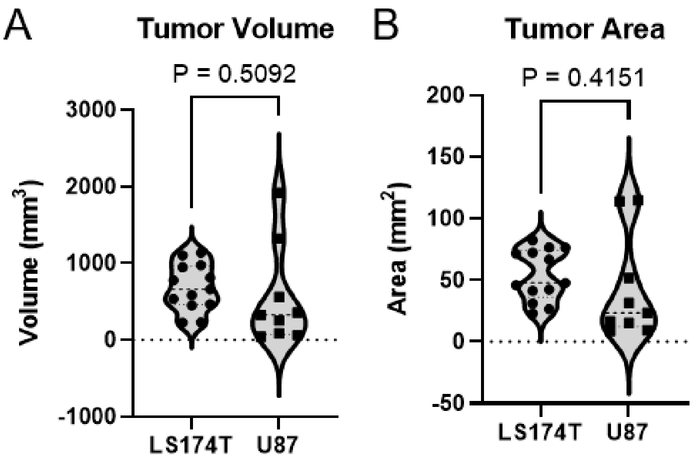


**Supplemental Fig. 2**. Tumor (A) volume and (B) area for LS174T and U87 tumors at the final ultrasound imaging time point before sacrifice.

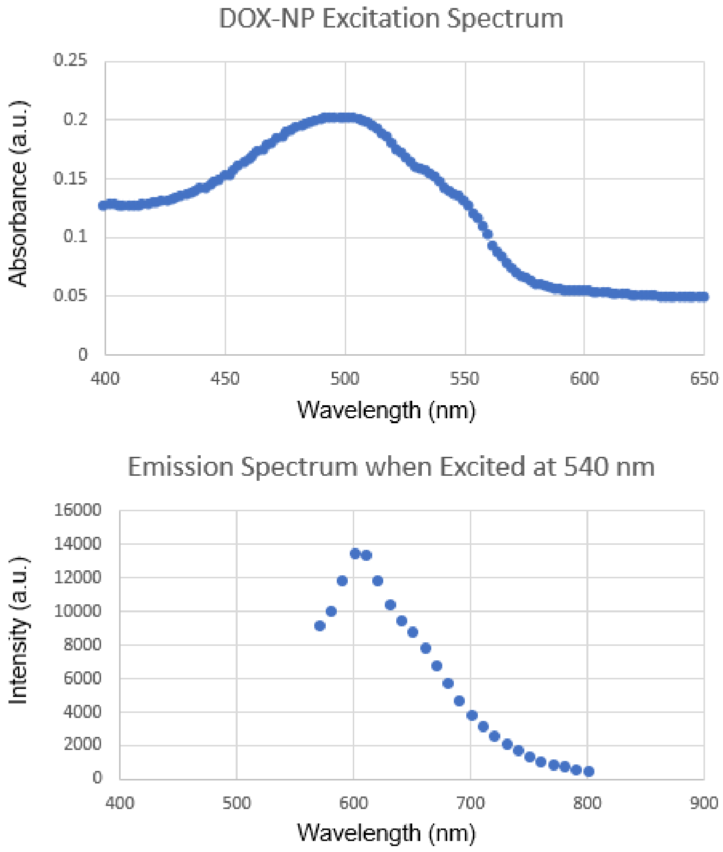


**Supplemental Fig. 3**. Excitation emission spectrum for doxorubicin-loaded liposomes via a TECAN fluorescent multimodal plate reader (Infinite M200, San Jose, CA).

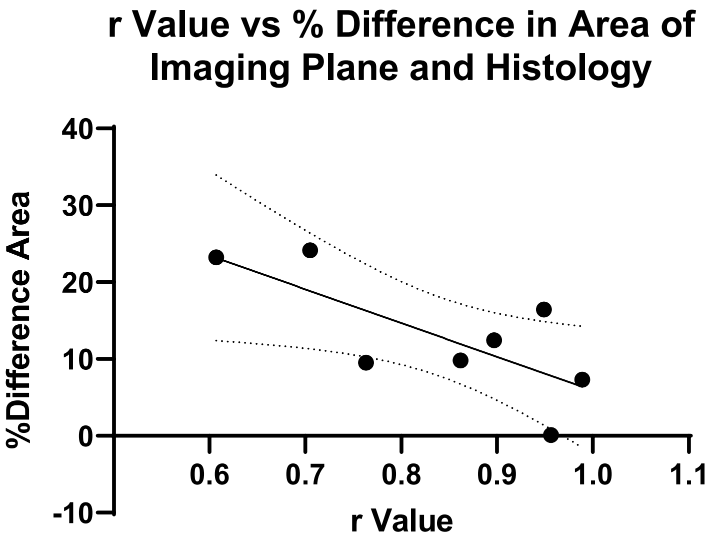


**Supplemental Fig. 4**. % difference between histology and ultrasound derived tumor area versus the intratumoral correlation coefficient found through quadrant analysis comparing DT to fluorescence intensity. This analysis uses all available LS174T data points.
